# Retinal stabilization reveals limited influence of extraretinal signals on heading tuning in the medial superior temporal area

**DOI:** 10.1101/643189

**Authors:** Tyler S Manning, Kenneth H Britten

## Abstract

Heading perception in primates depends heavily on visual optic-flow cues. Yet during self-motion, heading percepts remain stable even though smooth-pursuit eye movements often distort optic flow. Electrophysiological studies have identified visual areas in monkey cortex, including the dorsal medial superior temporal area (MSTd), that signal the true heading direction during pursuit. According to theoretical work, self-motion can be represented accurately by compensating for these distortions in two ways: via retinal mechanisms or via extraretinal efference-copy signals, which predict the sensory consequences of movement. Psychophysical evidence strongly supports the efference-copy hypothesis, but physiological evidence remains inconclusive. Neurons that signal the true heading direction during pursuit are found in visual areas of monkey cortex, including the dorsal medial superior temporal area (MSTd). Here we measured heading tuning in MSTd using a novel stimulus paradigm, in which we stabilize the optic-flow stimulus on the retina during pursuit. This approach isolates the effects on neuronal heading preferences of extraretinal signals, which remain active while the retinal stimulus is prevented from changing. Our results demonstrate a significant but small influence of extraretinal signals on the preferred heading directions of MSTd neurons. Under our stimulus conditions, which are rich in retinal cues, we find that retinal mechanisms dominate physiological corrections for pursuit eye movements, suggesting that extraretinal cues, such as predictive efference-copy mechanisms, have a limited role under naturalistic conditions.

**Significance Statement:** Sensory systems discount stimulation caused by the animal’s own behavior. For example, eye movements cause irrelevant retinal signals that could interfere with motion perception. The visual system compensates for such self-generated motion, but how this happens is unclear. Two theoretical possibilities are a purely visual calculation or one using an internal signal of eye movements to compensate for their effects. Such a signal can be isolated by experimentally stabilizing the image on a moving retina, but this approach has never been adopted to study motion physiology. Using this method, we find that eye-movement signals have little influence on neural activity in visual cortex, while feed-forward visual calculation has a strong effect and is likely important under real-world conditions.

## Introduction

In primates, optic flow is an indispensable cue for navigating through the world (Gibson, 1958; Warren and Hannon, 1988; Lappe et al., 1999). When the eyes are stationary, the brain can easily determine the instantaneous heading direction, which is the focus of expansion in the optic-flow pattern. When the eyes rotate during smooth pursuit, however, retinal slip distorts the optic flow and upsets this correspondence (Gibson, 1950). Despite these distortions on the retina, humans and non-human primates alike perceive very little distortion in their heading direction as they pursue moving objects (Royden et al., 1992; Britten and van Wezel, 2002). Such compensation is a general property of sensory systems, which often discount stimulation caused by the animal’s own behavior (termed reafferent stimulation). These results imply that the brain discounts distortions to the optic-flow field, but it is still unclear how this perceptual stability is maintained.

Two main classes of mechanisms by which the brain discounts the retinal-slip distortions have been proposed: *retinal* and *extraretinal* (Lappe et al., 1999; Britten, 2008). Under the retinal hypothesis, cortical areas selective for visual motion extract heading direction from the optic flow pattern directly by calculating and discounting motion components due to eye rotation (Royden, 1997; Perrone and Stone, 1998; Beyeler et al., 2016). Retinal mechanisms depend on depth cues to dissociate flow components due to retinal slip from those due to self-motion (Longuet-Higgins and Prazdny, 1980; Hildreth and Royden, 1998). On the other hand, the extraretinal hypothesis proposes that information about the retinal flow pattern is modified by an internal signal that tracks eye velocity to recover heading. These extraretinal signals likely originate from the efference copy (or corollary discharge) of motor commands for smooth pursuit rather than from proprioception (Sperry, 1950; von Holst and Mittelstaedt, 1950; Bridgeman and Stark, 1991; Crapse and Sommer, 2008). Psychophysical investigations have favored the extraretinal hypothesis, based on the comparison of perceived heading differences between normal pursuit and *simulated* pursuit, in which the eyes remain fixed while the experimenters artificially add rotation to the flow stimulus (Royden et al., 1994; Banks et al., 1996; Crowell and Andersen, 2001); but see (Warren and Hannon, 1988; Stone and Perrone, 1997; Li et al., 2006).

Neurophysiological evidence for retinal versus extraretinal mechanisms in self-motion processing is mixed. Neural responses in heading-selective areas like dorsal medial superior temporal area (MSTd) compensate for changes in the speed (Inaba et al., 2007; Chukoskie and Movshon, 2009) and direction of optic flow during smooth pursuit. Two classic studies reported a large extraretinal influence on the heading-direction preferences of MSTd neurons based on changes in activity between normal pursuit and simulated pursuit (Bradley et al., 1996; Shenoy et al., 2002). However, their stimulus lacked depth cues, which underlie many proposed mechanisms of retinally based corrections. Supporting this concern, MSTd neurons compensate much better for direction distortions when flow stimuli contain motion parallax and perspective cues to depth (Maciokas and Britten, 2010). However, that study could not identify whether the pursuit-invariant responses were due to a retinal or extraretinal mechanism.

In the present study, we designed a novel optic flow stimulus that isolates the effects of extraretinal influences on heading-selective neurons. The *stabilized pursuit* condition manipulates the relationship between eye rotation and the resulting retinal motion by rotating the stimulus with the eye as it pursues a target—effectively eliminating distortions to the optic flow while maintaining the influence of efference-copy signals. In this condition, as well as normal and simulated pursuit stimuli, we recorded from heading-selective MSTd neurons to identify the signal source responsible for the stability of heading responses during pursuit. We found that extraretinal mechanisms contribute only a small, though significant, amount to this stability, while retinal mechanisms have a considerably larger effect.

## Materials and Methods

### Animals and surgical procedures

Three adult female macaque monkeys (*Macaca mulatta*) were used in this study. Each monkey was surgically equipped with a head post, chronic recording cylinder (Crist Instrument Co.), and scleral search coil (Judge et al., 1980) to stabilize their heads, provide access for electrical recordings, and record their eye movements, respectively. Recording cylinders were placed under the guidance of prior structural MR images and stereotaxic atlases. All components were implanted under general anesthesia using sterile technique in a dedicated surgical suite. All procedures and experiments were performed in accordance with the National Institutes of Health guidelines and approved by the University of California, Davis Institutional Animal Care and Use Committee.

### Electrophysiological recordings

At the beginning of each recording session, we penetrated the dura mater with a stainless steel guide tube positioned within a polymer grid that ensured consistent access to the superior temporal sulcus. We then advanced single epoxy-coated tungsten microelectrodes (FHC) through the guide tube under the control of an electrical micromanipulator (National Instruments). Mulitunit activity was amplified (Bak Electronics), filtered for line noise, and passed through a dual voltage-time window discriminator (Bak Electronics) to isolate action potentials from single units. Timestamps from the individual spikes were then digitized at 1-ms intervals by the experimental control computer using the REX environment (Hays et al., 1982).

Before recording data for the main experiment, we mapped the dorsal subdivision of MST. Guided by MRI reconstruction and stereotaxic atlases, we identified MSTd based on the pattern of gray and white matter transitions as the electrode was advanced and previously described response characteristics (Tanaka and Saito, 1989; Graziano et al., 1994). Its neurons respond vigorously to patterns of moving dots, are often selective for complex motion patterns, and have large (compared to MT) receptive fields that often contain the fovea and portions of the ipsilateral visual field. We avoided recording from motion-selective cells in neighboring area 7a by ensuring that we recorded from a sufficiently ventral stereotaxic position.

### Visual Stimuli

We presented visual stimuli on a rear-projection screen with a PROPixx DLP LED projector (VPixx Technologies) at a display resolution of 1920 × 1080 pixels with a 120-Hz refresh rate. At 50 cm from the monkey, the projected image subtended 100°(horizontal) by 68°(vertical) of visual angle. The recording room was as dark as possible (minimum screen luminance of 0.78 cd/m^2^), and the monkey was kept in a light-adapted state by fully illuminating (110 cd/m^2^) the screen with white during the inter-trial intervals. These measures minimized the contributions of scattered light in the recording room to overall retinal motion. Throughout each experiment, we sampled eye position (National Instruments 12-bit ADC) at 1 kHz with a magnetic search coil system (DNI). We initially presented the stimuli under binocular viewing conditions for monkey Q, but we recorded the majority of the neuronal data under monocular occlusion of the ispilateral eye to reduce conflict between stereoscopic and motion parallax cues to depth.

To simulate self-motion in the main experiment, we developed a paradigm that translates and rotates a virtual camera through a 3D cloud of randomly positioned dots. During each trial, graphics commands were sent via a dedicated TCP/IP connection from the computer running the REX experimental control environment to a dedicated rendering machine. Stimuli were then generated on this machine with a custom software application that rendered each frame synchronously with the vertical refresh period of the projector.

In this environment, the viewable volume was a frustum (Figure 1A, top left) bounded by a near plane located at the surface of the screen, 50 cm from the observer, a far plane 150 cm from the observer, and the edges of the projected image on the screen. Dot density was 1000 dots/m^3^, which made roughly 3500 dots viewable at any time. Each dot was white (110 cd/m^2^) square that subtended ∼0.1° of visual angle on a black (0.78 cd/m^2^) background. No looming or changing disparity cues were present in this setup. The viewing frustum was embedded in a larger volume that ensured that the dot density was roughly constant as dots entered and exited the field of view. Throughout all self-motion conditions, translation speed was held at a constant 50 cm/s. Although the resulting pattern of dot motion on the screen is consistent with an infinite number of dot distance-observer speed combinations (if the ratio of the two is held constant), we refer to exact physical quantities here for clarity.

**Figure 1.**
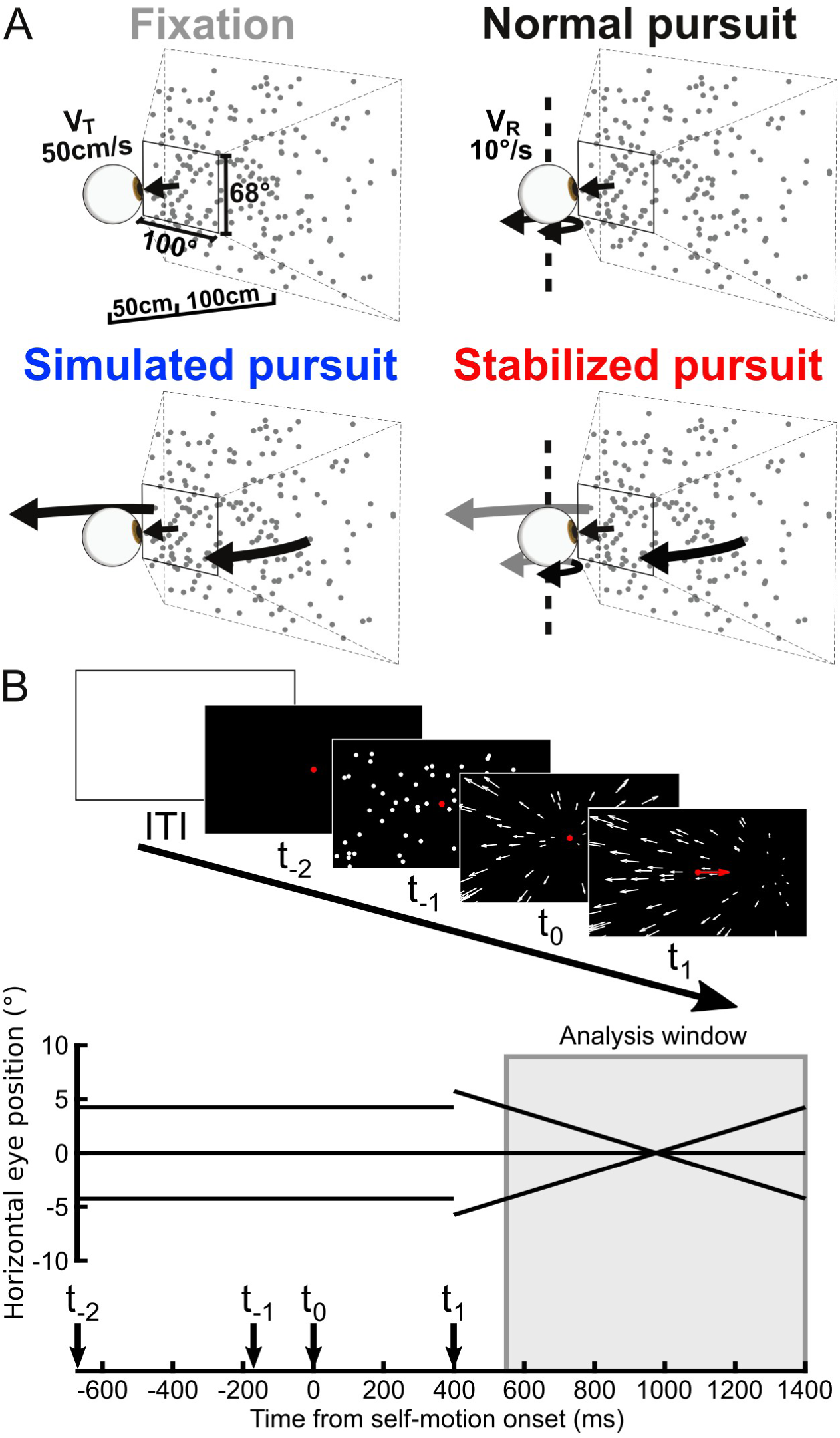
Experimental setup and trial time course. **A**, Geometry of the viewing volume and schematic of the three pursuit conditions. In all cases, heading was simulated by translating a virtual camera through a 3D space embedded with randomly placed dots. (Dot density was equal throughout the viewing volume.) For clarity, the contrast sign of the dots has been inverted from what was presented. The dots have also been increased in size and decreased in number. In the normal and stabilized pursuit conditions, the red target moved to the right or left at 10°/ s, independent of the background dots. In simulated pursuit, the red target remained fixed while the virtual camera was rotated right or left. For the stabilized pursuit condition, the virtual camera was rotated with the same rotational velocity of eye while the monkeys pursued the target. **B**, Trial time course and eye position traces. Before and after each trial begins (ITI), the screen is fully illuminated. After the red fixation dot appears on the screen (t_-2_) at one of three locations, the monkey must remain fixated for 500ms before the dots appear (t_-1_) in the volume. Following a 180-ms pause, the translation epoch (t_0_) begins as the monkey continues to fixate. For pursuit trials, the fixation dot initially steps at the beginning of the epoch (t_1_) to a more eccentric position before traveling to the left or to the right.

Depending on the stimulus condition, the monkey either fixated on a central dot or pursued a moving target during simulated self-motion. This target was a red dot that subtended 0.25° of visual angle and moved independently of the other dots embedded in the 3D environment. Each monkey had to remain within a 1.75° square window during fixation or a 2° window during pursuit; otherwise the trial was aborted. To minimize the number of catch-up saccades, the pursuit target on all but the earliest experiments moved in a step-ramp fashion (Rashbass, 1961), with the initial step magnitude (Figure 1B) held constant across monkeys and chosen to roughly approximate the lag time between target motion onset and smooth pursuit initiation.

### Pursuit manipulations

The stimuli in the main experiment consisted of four different pursuit conditions over the same set of simulated heading directions (Figure 1A). In the first condition, fixation, the monkey simply had to remain fixated on a central target while we simulated self-motion. The second condition, normal pursuit, required the animal to pursue the target moving either leftward or rightward in the plane of the screen during simulated self-motion. In the next two conditions, the correspondence between eye rotation and the resulting reafferent motion on the retina was manipulated. During simulated pursuit, the monkey remained fixated on a central target while the effects of reafferent motion on optic flow were simulated by rightward or leftward rotation of the camera as it translated through the virtual environment. This produced a dynamic retinal image identical to that found in normal pursuit while the eyes were stationary. Finally, in our stabilized pursuit condition, we eliminated reafferent motion while the eyes were in motion by using online estimates of instantaneous eye velocity to rotate the camera in the same direction. This produced a retinal flow pattern nearly identical to that found during fixation.

To estimate instantaneous eye velocity for stabilization, we first took a 10-ms sliding window average of eye position throughout the trial. From this running average, eye velocity was estimated with numerical differentiation and used to estimate the rotation of the camera view between frame draws. This rotation was incorporated into the calculations used for the simulated translation of the camera for the next frame (Figure 2). Across all trials, this resulted in roughly a single-frame lag (mean = 9.6 ms) between changes in eye velocity and the resulting corrections on the screen (Figure 2, inset).

**Figure 2.**
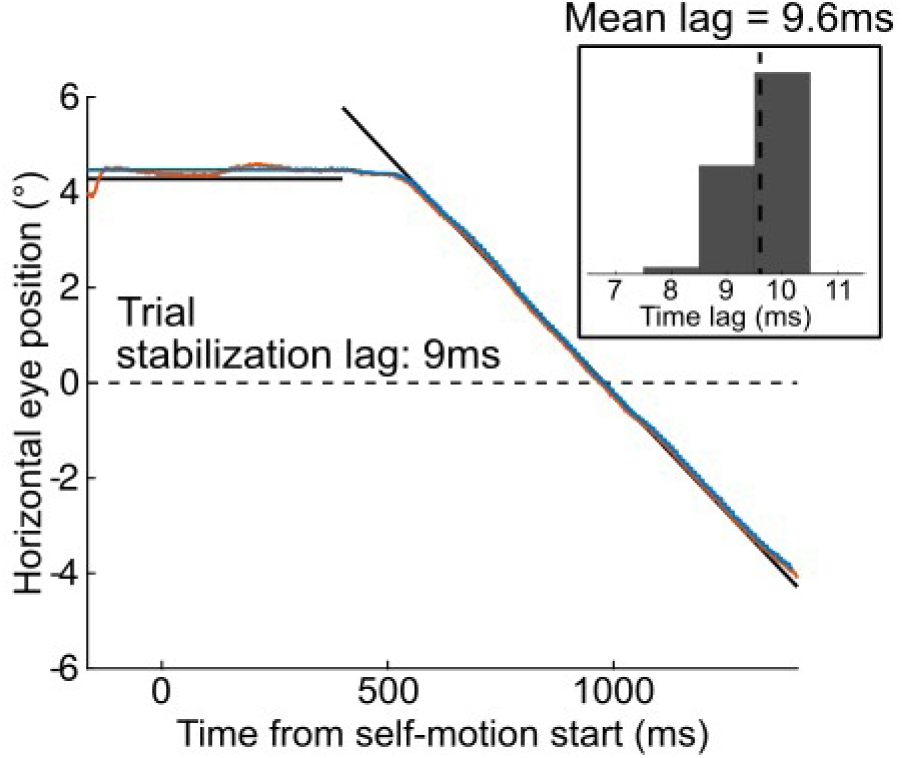
Performance of retinal stabilization in a single trial and across all stabilized pursuit trials. In the trial shown, the pursuit target (black) is moving to the left at 10°/s. Eye position is shown in orange while the corrective rotations to the camera viewing angle are shown in blue. For each trial, stabilization lag was determined by finding the lag time that maximized the cross-correlation between the eye position and camera position traces. **Inset**, histogram of stabilization lag times across all stabilized pursuit trials. Mean lag time was calculated to be 9.6 ms, which is roughly a single frame lag with the projector refresh rate at 120 Hz.

### Experimental protocol

Upon isolation, each cell was initially characterized for its heading and pursuit direction preferences. Heading preference was estimated by simulating self-motion in 26 evenly spaced directions in 2D heading space (i.e. elevation and azimuth) while we recorded spiking activity from a given cell. To reduce the number of unique conditions to a manageable number, we used these responses to select a subset of headings for presentation in the main experiment. In this subset, we chose heading directions at single elevation that were centered around the heading that evoked the maximal response from each neuron. Heading azimuths were chosen to span a range that covered most or all of the cell’s response range, encompassing the peak response when possible. Cells were also hand-mapped to estimate the spatial extent of their receptive fields and their rough tuning in spiral space (Graziano et al., 1994). When cells could be held long enough, we ran an additional automated procedure to measure their tuning to planar and spiral space motion after the main experiment. Pursuit direction preferences were estimated by presenting targets moving in 8 equally spaced directions in the plane of the screen without optic flow stimuli. We also included a target blink period in the middle of the ramp epoch to assess cells for tuning to extraretinal signals related to pursuit (Newsome et al., 1988). During this brief (150 ms) period, the pursuit target was extinguished (i.e., no stimuli were present in the animals’ visual field) while monkeys continued to pursue the implied path of the target.

During the main experiment, the four pursuit conditions were pseudorandomly interleaved. Each trial (Figure 1B) consisted of an initial fixation epoch followed by a brief period in which the dot-filled viewing volume appeared before the onset of camera translation. Once the virtual camera started moving, the pursuit target remained stationary for 400 ms before following the step-ramp trajectory to the left or right. Neuronal activity during pursuit initiation was ignored, and spikes were counted during a window (Figure 1B, shaded box) for subsequent analyses. To control for the effects of eye position on neural responses (Bremmer et al., 1997), we positioned the pursuit target such that it had the same mean position across the length of this analysis window in each of the pursuit conditions (Figure 1B, eye position traces). Finally, each trial was followed by an intertrial interval in which the screen was fully illuminated to maintain light adaptation. This eliminated retinal slip caused by other objects in the recording room that were dimly illuminated by light scattering off the screen.

### Effects of pursuit on retinal flow patterns

In order to estimate the magnitude of retinally based compensation for pursuit in MSTd, we developed a method to compare a neuron’s heading preferences during simulated pursuit to an estimate of the cell’s heading preference if it did not compensate at all. We determined this non-compensating response by analyzing the distortions to optic flow during smooth pursuit. In a single plane, retinal slip from pursuit shifts the center of motion of an expanding pattern in the direction of eye movement (or vice versa for contracting patterns). For visual scenes with more than one depth plane, the magnitude of these shifts increases with distance from the observer, producing an apparent curvature in flow pattern on the retina. We reasoned that a MSTd neuron that did not compensate for pursuit would match a heading direction to these distorted patterns by aggregating center of motion shifts across depth planes into a single location. Therefore, we can estimate how much this non-compensating neuron would appear to shift its heading preference by subtracting this matched heading direction from the neuron’s preferred heading direction.

To model the expected effects of smooth pursuit on neuronal responses, we derived a set of differential equations to describe the motion of a texture element for an arbitrary set of heading and eye rotations. To do so, we followed the method of Longuet-Higgins and Prazdny (1980) while incorporating the screen distance of our setup into the perspective projection (i.e. *x = D*_*screen*_*X/Z*):

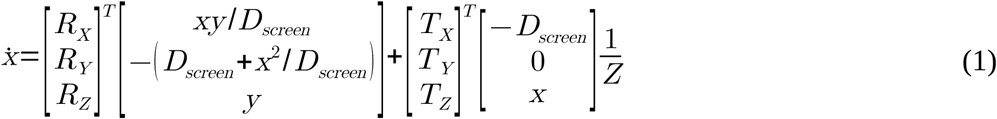

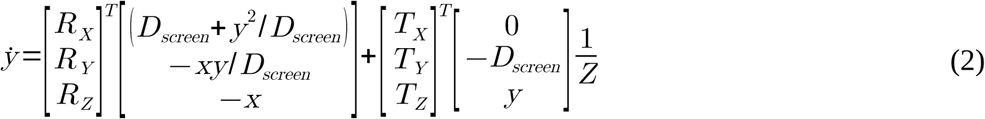

Where *X, Y*, and *Z* are the positions of dots in 3D space, *x* and *y* are the horizontal and vertical screen positions of each projected dot, *R*_*X*_, *R*_*Y*_, and *R*_*Z*_ are the components of the eye’s rotation about each respective axis, *T*_*X*_, *T*_*Y*_, and *T*_*Z*_ are the translational components, and *D*_*screen*_ is the distance of the screen from the observer. If we assume that the center of motion determines each MSTd neuron’s response to different retinal flow patterns, and restrict eye rotations to those about the Y axis, as we do in this experiment, we can calculate the horizontal (*x*_*CoM*_) and vertical (*y*_*CoM*_) position of this center on the screen as follows:

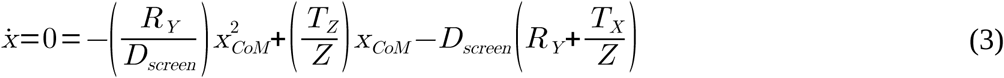

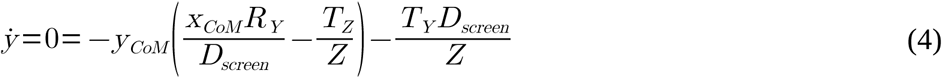

Solving for the quadratic in (3) yields the horizontal component of the center of motion for any arbitrary combination of heading direction and eye rotation. Additionally, we find that the x component of this center is independent of the *T*_*Y*_ component of the heading direction, so we can determine the expected horizontal shift in center of motion in the presence of left or right pursuit for any given heading direction. To retrieve the vertical component, we can rearrange (4) to produce:

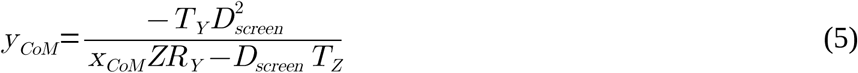

Here, we find that there will be no vertical shift in the center of motion during left or right pursuit for any heading with *T*_*Y*_ = 0 (i.e., self-motion parallel to the ground plane). Finally, the following transform expresses the position of the center of motion on the tangent screen in degrees of visual angle:

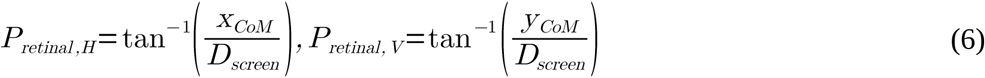

These equations were used to estimate the magnitude of tuning shifts in a hypothetical neuron that did not compensate for retinal flow distortions during pursuit. We first determined the shift in the center of motion for each stimulus depth plane in 1-cm increments of depth. From this, we determined the purely translational heading direction that best matched the aggregate center of motion in the pursuit-distorted retinal flow pattern. To do so, we calculated the mean center of motion over the nearest visible 10 cm of the stimulus (50-59 cm), or just under 1% of the total volume of the viewing frustum. This value was chosen as a conservative estimate of the neurons’ weighting of each depth plane (as the nearest planes shift less than the farthest ones) while also allowing enough dot motion (∼35 dots) to be visible for a reasonable amount of time (∼20-200 ms). Using either an inverse depth-weighted mean of the set of centers of motion (to reflect velocity scaling due to motion parallax) or an unweighted mean produced larger, but roughly similar results. We then calculated predicted tuning shifts using a numerical method (with MATLAB’s fmincon function) that found the heading direction that minimized the difference between the aggregate center of motion position and the center of motion position that matched the cell’s heading preference.

### Data analysis and statistical testing

All eye position and electrophysiological data were analyzed using custom scripts in MATLAB (Mathworks, RRID:SCR_001622). We determined the lag between changes in eye velocity and subsequent corrections on screen during the stabilized pursuit trials by finding the lag time that maximized the cross-correlation between the camera and eye-position traces for each trial (Figure 2). Before the cross-correlation, the camera position traces were upsampled from 120 to 1000Hz. Trial-wise lags were then aggregated, revealing a mean stabilization lag time of 9.6 ms.

To quantify how well the monkey followed the pursuit target, we calculated the pursuit gain and number of saccades for each trial. We estimated instantaneous eye velocity (in °/s) with numerical differentiation of the raw eye position (*p*) using a symmetric difference quotient:

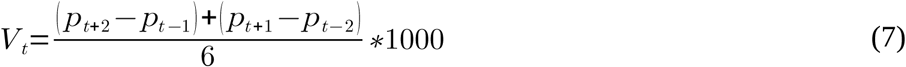

Differentiation was followed by smoothing with a 10-ms sliding-window average. We calculated pursuit gain by dividing the average pursuit velocity in the analysis window by the constant velocity of the pursuit target. Saccade episodes were identified by finding whenever the estimated 2D eye velocity exceeded the standard deviation by a factor of three for that trial.

Cells were selected for inclusion based on four criteria: isolation quality, minimum number of trial repeats, significant heading tuning, and Gaussian-like tuning for heading direction. The spike trains of each cell had to exhibit a clear refractory period as assessed by autocorrelation. To be included, each unit had to have at least four repeats for each of the 49 conditions in the main experiment (mean number of repeats per condition was 8). As a first pass to ensure we included only heading-selective neurons, we used a one-way Welch’s ANOVA test for significant firing differences between the 26 presented directions in the heading tuning protocol. Units that passed this test had their response patterns under the fixation condition fit with the following von Mises function using MATLAB’s fmincon:

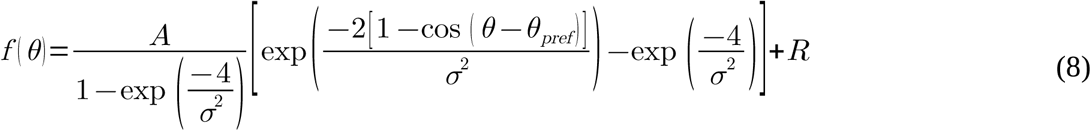

Where *A* is an amplitude parameter, *θ* and *θ*_*pref*_ are the presented and preferred heading directions, *σ* is a parameter that sets heading selectivity, and *R* is the response offset that sets the response of the cell to the antipreferred heading (Figure 4A, bottom). Upper and lower bounds for the parameters were chosen to be physiologically realistic (hard lower bound at 0 for amplitude and response offset), to avoid excessively peaked fits (σ ≥ 1.5 times inter-heading spacing), and to keep the estimate of preferred tuning within one inter-heading spacing of the most extreme angles tested. Cells with poor fits during the fixation condition (r^2^<0.5) were excluded from all further analysis.

**Figure 4.**
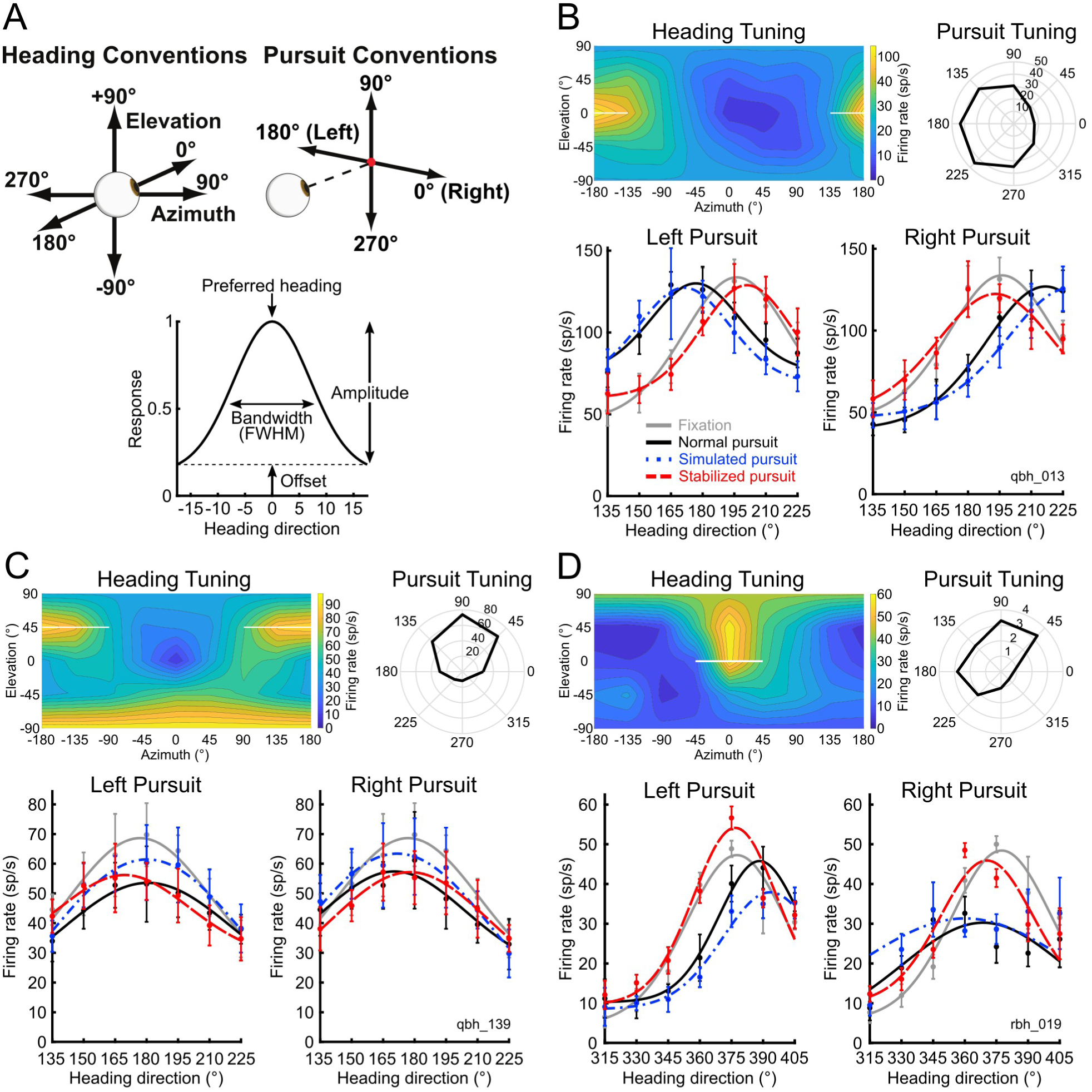
Example MSTd unit responses. **A**, Top, heading and pursuit direction conventions. Bottom, naming conventions for cell response parameters extracted with von Mises curve fit. **B-C,** Examples of single-unit responses under pursuit manipulations. Heading tuning is shown with an equirectangular projection of 2D heading space. White bars in heading tuning plots denote the range of headings tested in the main experiment. Error bars represent standard deviation in firing rate across the repeats of each condition. **B**, A unit that is broadly tuned for leftward pursuit and backward heading directions. **C**, A unit tuned for upward pursuit and broadly tuned for backward heading. This unit exhibited substantial tuning for clockwise spiral motion. **D**, An example unit that is suppressed during the smooth pursuit tuning protocol (mean response during fixation: 6 sp/s). The cell is also well tuned for forward and slightly upward heading direction.

In the main experiment, the responses of each cell in each of the six pursuit conditions (normal pursuit, simulated pursuit, and stabilized pursuit to the left or right) to the seven presented heading directions were fit individually with the von Mises with all four parameters free. To obtain the tuning bandwidth (full-width of tuning curve at half-maximum) from these fits, we used the following equation:

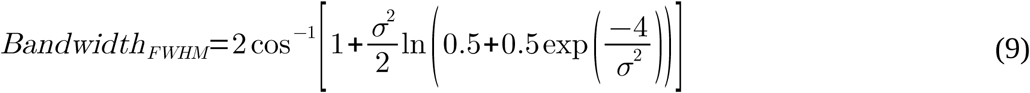

Since our analyses used pairwise comparisons of parameter fits between fixation and pursuit, if any pair member contained a fit that was not sufficiently better than a mean fit (threshold r^2^>0.5), the pair was excluded from further analysis.

To combine changes in preferred heading direction *θ*_*pref*_ between leftward and rightward pursuit trials as well as in cells that preferred forward and backward headings, we needed to take into account that these conditions induce different directions of curvature in the retinal flow pattern. For example, rightward pursuit produces rightward curvature for forward heading, but leftward curvature for backward headings (and vice versa). Because these distortions should produce different signs of tuning curve shifts between the fixation and pursuit conditions, we reversed the signs for data obtained under leftward pursuit and backward heading so that all analyzed shifts are consistent with rightward pursuit and forward heading.

To test for significant tuning shifts in the cell sample, we used a Wilcoxon signed-rank test to assess the paired difference between heading preferences during fixation and during each of the three pursuit conditions after the sign-reversing procedure. We used the same test to compare heading preference shifts between normal and simulated pursuit (Figure 8), and between monocular and binocular viewing conditions (Figure 10).

**Figure 8.**
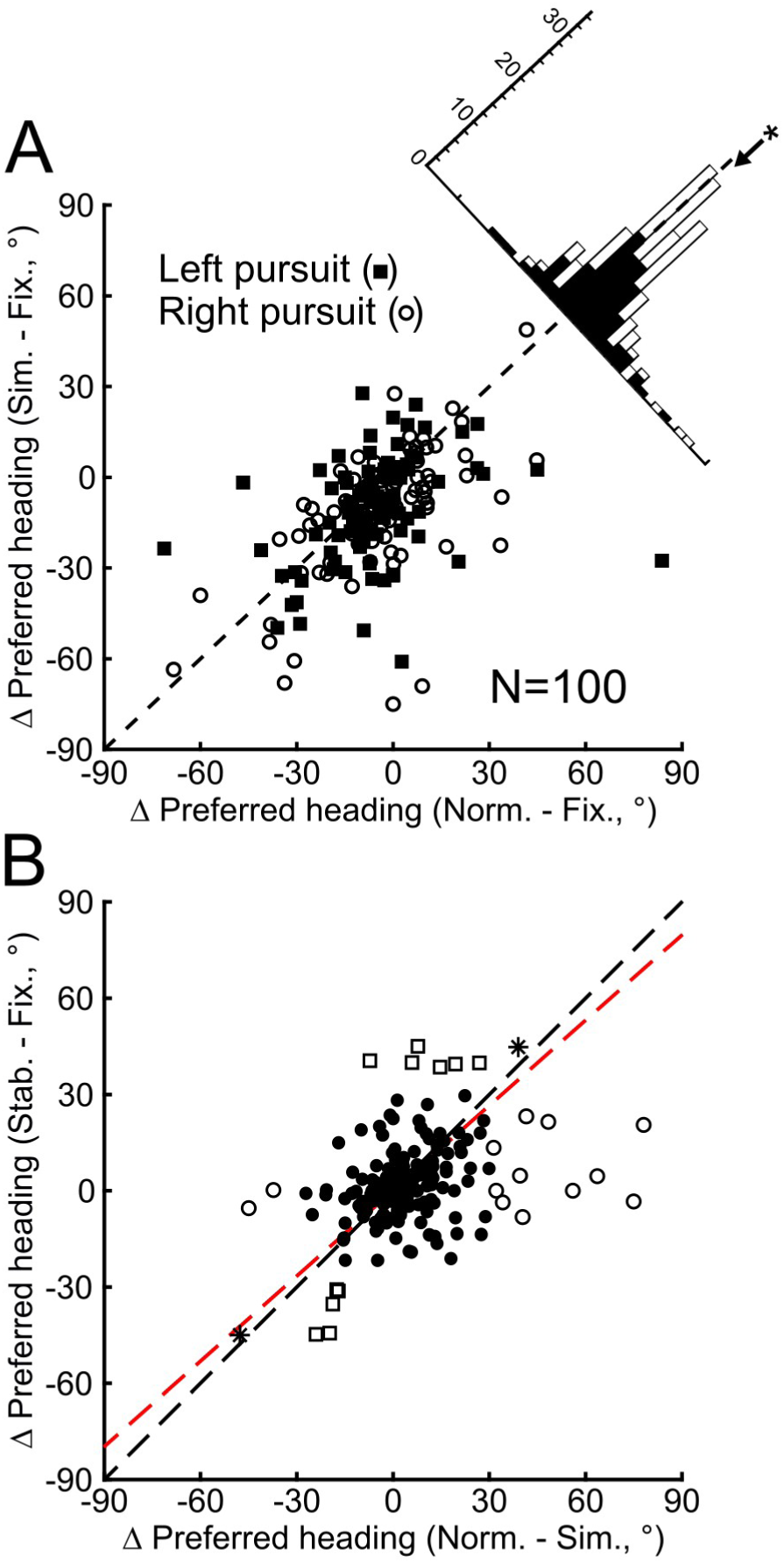
Effects of efference copy on heading tuning in cell sample. **A**, Scatter plot and marginal histogram comparing the changes in heading-direction preferences between the simulated and normal pursuit conditions and fixation for all included units. **B**, Paired comparison of heading shifts for each unit between normal and simulated pursuit and between the stabilized pursuit and fixation conditions. Open squares denote significant stabilized pursuit vs. fixation differences, open circles denote significant normal vs. simulated pursuit differences, and asterisks denote significant differences for both comparisons.

**Figure 10.**
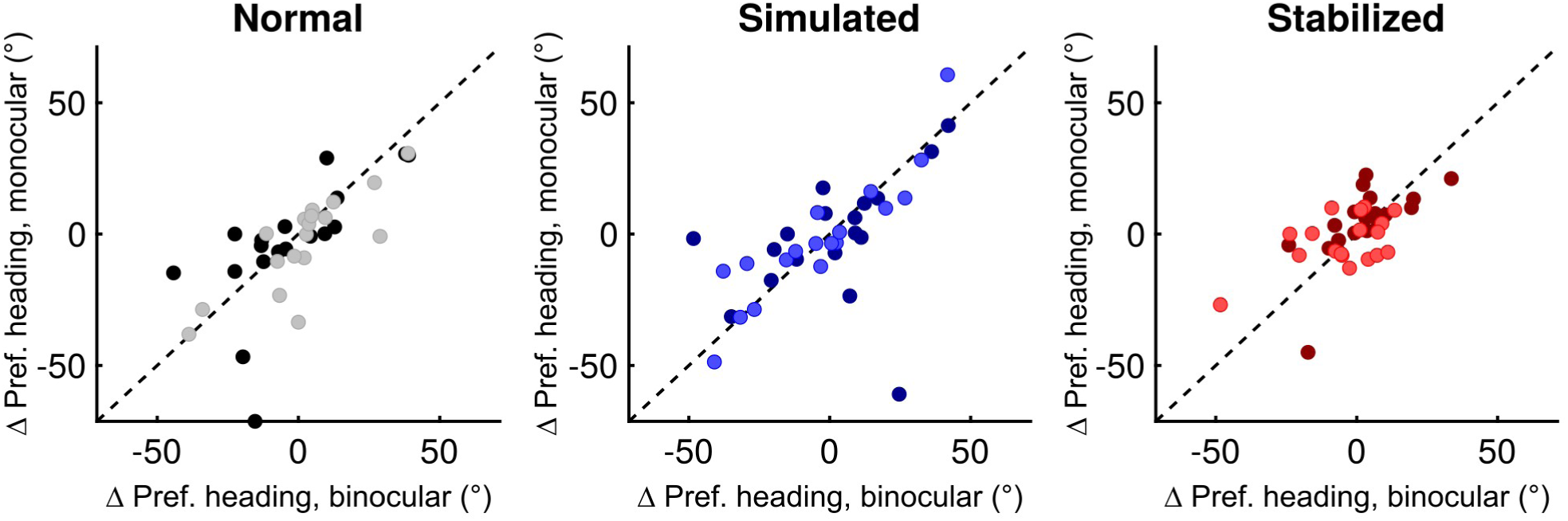
Comparison of heading preference shifts between monocular and binocular viewing conditions. Unit responses (N = 19) under each of the pursuit conditions were recorded with and without monocular occlusion; the starting viewing condition was pseudorandomized between cells. Dashed black line represents line of unity. Pursuit direction color conventions are the same as for Figure 6.

For the amplitude and bandwidth parameters, we assessed differences between fixation and pursuit with a Wilcoxon signed-rank test in a similar manner. Given the distribution of fits, we tested for pairwise differences in the offset parameter using the following permutation procedure. For each pursuit condition, we pooled the calculated offset parameters from both fixation and pursuit and selected values (without replacement) to form two new samples and calculated the median paired difference between them. This procedure was then repeated 10000 times to obtain a distribution under the null hypothesis, which was then used to calculate the probability of obtaining the actual median paired difference.

A model II regression was used to quantify the paired relationship between the simulated-normal pursuit and stabilized pursuit-fixation heading preference shifts (Figure 8B). The best fitting line was estimated with the line bisecting the minor angle between the Y-on-X and X-on-Y regression lines, which is robust to differences in range between the two measurements (Sprent and Dolby, 1980). In addition, we calculated the Spearman’s rank correlation coefficient between the two pairs of differences and assessed the significance of this result with a t-test. To identify neurons that showed significant differences between stabilized pursuit and fixation and between normal and simulated pursuit, we used a permutation test to construct a distribution under the null hypothesis. Responses from individual trials were pooled within each of these pairs (separately for each heading direction) and used to generate new sets of responses for each of the 49 different conditions. These shuffled responses were again fit with a von Mises function to estimate heading preference shifts. After repeating this procedure 500 times, tuning shifts in the original data set were deemed significant with a two-tailed test at the 5% level.

To compare the differences between expected tuning shifts and measured shifts during simulated pursuit, we used a Model I linear regression and a Wilcoxon Signed-Rank test for paired differences.

### Code accessibility

All data analysis and modeling code used to define uncorrected heading preference shifts can be accessed as freeware from https://github.com/tsmanning/EfferenceCopyMST.

### Software accessibility

The software developed for stimulus presentation and image stabilization (*render*) is available upon request.

## Results

### Manipulating the relationship between eye velocity and retinal slip isolates the effects of retinal and extraretinal signals on neuronal responses to optic flow

To investigate the origin of the corrective signals for pursuit in MSTd, we manipulated the relationship between retinal slip and rotational eye velocity as monkeys viewed a set of stimuli simulating self-motion through a 3D scene filled with randomly placed dots (Figure 1A). In our unmanipulated baseline condition, normal pursuit, each monkey pursued a red target that moved to the left or the right independently of the dots in the optic flow pattern. In the simulated-pursuit condition, we recreated the same retinal flow pattern present during normal pursuit by rotating the viewing direction of the virtual camera while the monkey fixated on a stationary central target (Figure 1A, bottom left). Because the eyes were stationary, this condition eliminated efference-copy inputs and therefore isolated the effects of retinal mechanisms of flow stabilization on MSTd activity.

We also developed a novel manipulation that largely eliminated the distortions to the pattern of retinal flow arising from pursuit and therefore isolated the contributions of extraretinal mechanisms. In this stabilized-pursuit condition (Figure 1A, bottom right), the monkey pursued a moving target as in normal pursuit, but we rotated the virtual camera in the same direction as the eye rotation based on online estimates of eye velocity (see Methods). As a result, the pattern of retinal flow was nearly identical to the undistorted pattern present during the fixation condition, even though the monkey still pursued the moving target. To ensure that stabilization occurred with the shortest lag possible, we updated our stimulus at a high frame rate and estimated eye velocity online with scleral search coils. The corrective rotations to the virtual camera faithfully matched those of the eye during pursuit with a mean lag roughly equal to a single frame update cycle (9.6 ms, Figure 2).

Pursuit eye movements distort optic flow by introducing an apparent curvature in depth to the motion pattern in the same direction as the eye rotation for forward heading directions (Figure 3A, compare upper and lower flow patterns). More specifically, rightward pursuit produces rightward shifts in the centers of motion that increase proportionally with the distance from the viewer. Therefore one would expect a substantial shift in the tuning curve if MSTd neurons acted as eye-centered (Fetsch et al., 2007; Lee et al., 2011) flow filters, reflecting the degree to which the center of motion in the stimulus overlapped with the center of motion associated with the cell’s preferred heading direction. However, pursuit eye movements produce only small shifts in the tuning curves of heading-selective neurons in MSTd opposite that of the center of motion shifts (Figure 3A, bottom) (Bradley et al., 1996; Shenoy et al., 2002; Maciokas and Britten, 2010). These shifts reflect an under-compensation for pursuit-related distortions. If the responses of MSTd neurons were completely stable during pursuit, the tuning curves would completely overlap. On the other hand, one would expect a substantial shift in the tuning curve if MSTd neurons acted as eye-centered (Fetsch et al., 2007; Lee et al., 2011) flow filters, reflecting the degree to which the center of motion in the stimulus overlapped with the center of motion associated with the cell’s preferred heading direction. For example, the flow pattern resulting from translational self-motion and eye rotation in Figure 3A that best matches the preferred pattern of the neuron without pursuit is displaced by ∼30 degrees to the left, which should produce a similar shift in the tuning curve. As the difference between the peaks of the curves is much closer to 0° than 30°, we can surmise that MSTd uses either efference copy or a retinal cue to pursuit eye movements to achieve stability.

**Figure 3.**
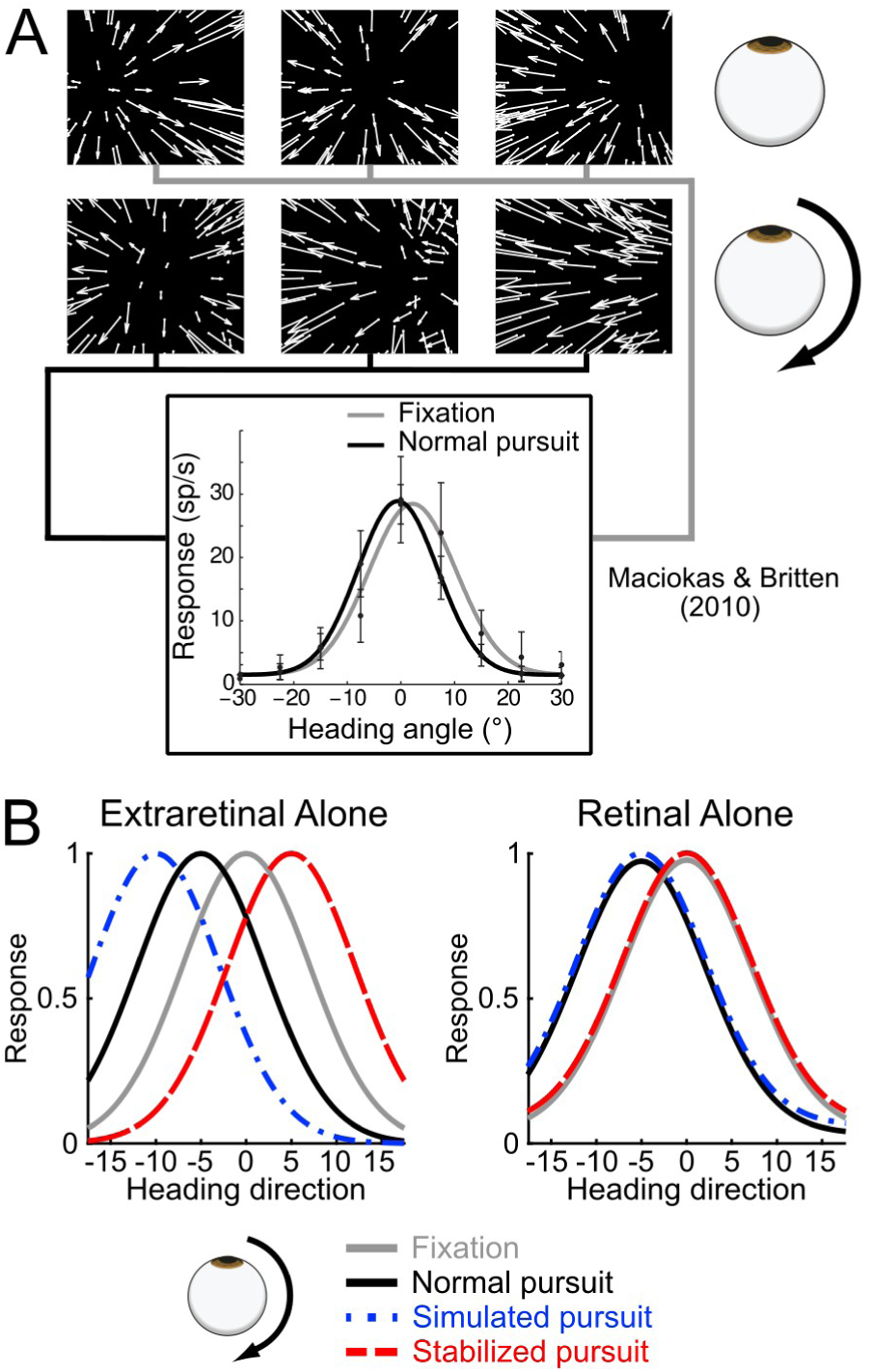
Effects of smooth pursuit on retinal flow and neuronal responses, and predictions under two hypothesized mechanisms of heading stability. **A, Top**, optic flow patterns for three different heading directions about straight ahead (−30°, 0°, +30°). **Middle**, retinal flow patterns for the same three heading directions during rightward pursuit. Summing the motion components due to self-motion and eye rotation produces an apparent rightward curvature to the flow pattern due to increasing rightward shifts in the centers of motion with each successive depth plane. **Bottom**, results from Maciokas & Britten (2010) demonstrating the effects of these distortions on the responses of a single unit in MSTd. Given the leftward displacement of the center of motion in each of the flow patterns with rightward pursuit, the cell’s tuning curve is expected also to move substantially to the left; yet it shifts only a small amount. **B**, predicted tuning curve shifts under each of the pursuit conditions in the present study in the case where cells achieve stability for heading direction encoding during pursuit using purely a retinal or purely an extraretinal signal.

If we assume that efference-copy and retinal-cue mechanisms are mutually exclusive, we can predict how a neuron’s heading-direction preference will change under each stimulus condition. Under the hypothesis that efference copy alone is responsible for response stability, we would expect to observe the responses at the left of Figure 3B during rightward pursuit from a neuron that prefers forward self-motion (0°Az). Responses during fixation and normal pursuit would be similar to those at the bottom of Figure 3A, with a small leftward shift in the tuning curve during pursuit. In simulated pursuit, we eliminate the influence of efference copy on MSTd neurons while retaining the distorted pattern of retinal flow. We therefore would expect to see an even greater leftward shift, indicating that the cell signals the true shift in the flow pattern’s center of motion on the retina. Conversely, in stabilized pursuit, we retain the putative efference copy inputs while eliminating the flow distortions. If we assume efference copy pushes the preferred heading direction of the cells in the direction of pursuit to counteract the distortions (i.e. shifts the simulated pursuit tuning curve toward the normal pursuit curve), we would expect to see a rightward shift in tuning compared to fixation during stabilized pursuit. We therefore would expect the difference in tuning between simulated and normal pursuit and between fixation and stabilized pursuit to be of the same magnitude and opposite sign, as efference copy should have the same effect in both cases.

Under the hypothesis that retinal mechanisms alone account for response stability, we would predict the responses at the right of Figure 3B. In this case, we assume that MSTd neuronal responses will be purely determined by the retinal flow pattern. As shown previously, during normal pursuit we would see a leftward shift in tuning compared to fixation. The magnitude of this shift is assumption-dependent; we model this quantitatively for each cell below. In simulated pursuit, we would expect the cell’s responses to be identical to those found in normal pursuit, as the distorted flow patterns are identical in the two conditions. Likewise, motion on the retina is identical and undistorted under both fixation and stabilized-pursuit conditions, so we would expect neuronal responses to be identical in these conditions.

### Pursuit manipulations alter preferred heading directions in MSTd neurons

To test these predictions, we recorded from MSTd neurons in three female macaque monkeys (Monkey P: left hemisphere, Q: right hemisphere, R: left hemisphere) while they performed the fixation or pursuit tasks. Before the main experiment, we categorized each cell in terms of heading, spiral space, and pursuit tuning (see Methods). To estimate the changes in preferred heading direction between the conditions, we fit the set of responses in each condition independently with a modified von Mises function (see Methods). Each parameter of the tuning curve (Figure 4A, bottom) was free to vary during the fitting procedure. Three example cells (Figure 4B-D) illustrate that retinal mechanisms are responsible for the majority of compensation for pursuit eye movements based on the predictions outlined in Figure 3B.

We recorded full data sets from 127 well-isolated neurons, of which 101 cells passed our conservative inclusion criteria; we excluded cells that were non-selective (p > 0.05 for Welch’s ANOVA) or had response patterns during fixation that were poorly fit with a von Mises function (r^2^ < 0.5). For each included cell, we also excluded stimulus conditions (i.e., pursuit direction or manipulation condition) in which responses were poorly fit (Table 1). The heading preferences were biased for forward self-motion (Figure 5), characteristic of MSTd (Graziano et al., 1994; Takahashi et al., 2007). Pursuit preferences were fairly evenly spread across the eight pursuit directions tested.

**Table 1.** Number of included cells in each pursuit condition.

|  | Monkey P (L,R) | Monkey Q (L,R) | Monkey R (L,R) | Total (L,R) |
| --- | --- | --- | --- | --- |
| Fixation | 15 | 70 | 16 | 101 |
| Normal pursuit | 15 (14,14) | 70 (68,67) | 16 (16,15) | 101 (98,96) |
| Simulated pursuit | 15 (14,14) | 70 (68,68) | 16 (16,14) | 101 (98,96) |
| Stabilized pursuit | 15 (15,13) | 70 (67,67) | 15 (15,12) | 100 (97,92) |

Number of cells that passed the inclusion criteria independently determined for each pursuit condition (left or right). Total number of unique cells for each monkey precede the individual numbers for each of the conditions in parentheses. The total count may be larger than either of the subtotals for left or right pursuit if the cell sets are not identical (i.e., the results from one cell passed inclusion for left pursuit but not for right pursuit or vice versa).

**Figure 5.**
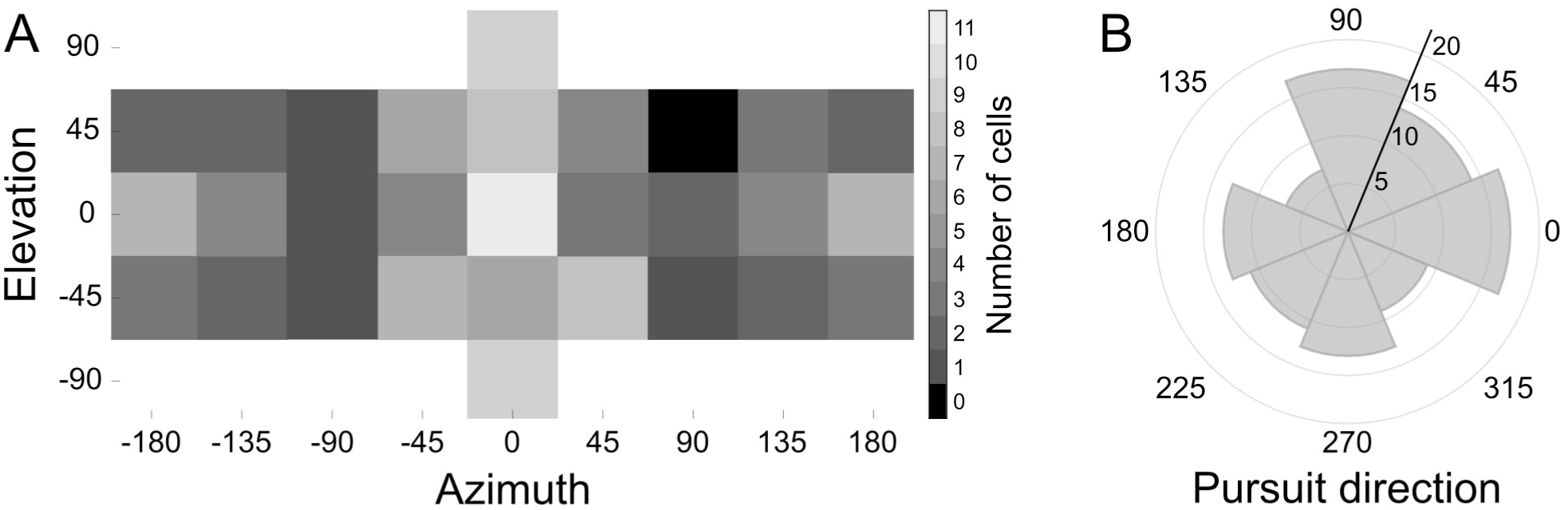
Distribution of heading and pursuit direction preferences for all included cells in the sample. **A**, Histogram of heading preferences determined by direction of maximal response. **B**, Polar histogram of pursuit preferences as determined by direction of maximal cell response.

We initially recorded neurons under binocular conditions but recorded the bulk of the data under monocular occlusion to minimize cue conflict between motion parallax and binocular disparity cues to depth. To compare the effects of the two conditions, we also ran the main experiment under both monocular and binocular viewing on a subset of cells. For cells recorded under both viewing conditions, only the monocular data were included in the subsequent analyses (Table 2).

**Table 2.** Number of included cells under each viewing condition.

|  | Monkey P | Monkey Q | Monkey R | Total |
| --- | --- | --- | --- | --- |
| Monocular | 15 | 41 | 13 | 69 |
| Binocular | 0 | 29 | 3 | 32 |
| Total | 15 | 70 | 16 | 101 |

Number of cells included in the analyses for each monkey under each viewing condition (monocular or binocular, M or B).

To analyze the contribution of retinal and extraretinal mechanisms to pursuit compensation, we compared fitted heading preference between fixation and each of the pursuit conditions (Figure 6). Because optic flow patterns are opposite for forward (expanding patterns) and backward (contracting patterns) headings, pursuit in a given direction will produce opposite shifts in the centers of motion and tuning curve peaks. Similarly, leftward and rightward pursuit directions produce opposing directions of retinal slip and therefore shifts in the centers of motion (Figure 6A). For forward headings (unshaded regions of plot) and rightward pursuit, the dots scatter below the line of unity, consistent with an overall counterclockwise shift in heading preferences across the cell sample. This pattern is reversed for backward headings (shaded regions), and the overall pattern of preference shifts is reversed for leftward pursuit (compare locations of light and dark dots). Overall, these results corroborate the slight under-compensation for pursuit seen in the single cell examples.

**Figure 6.**
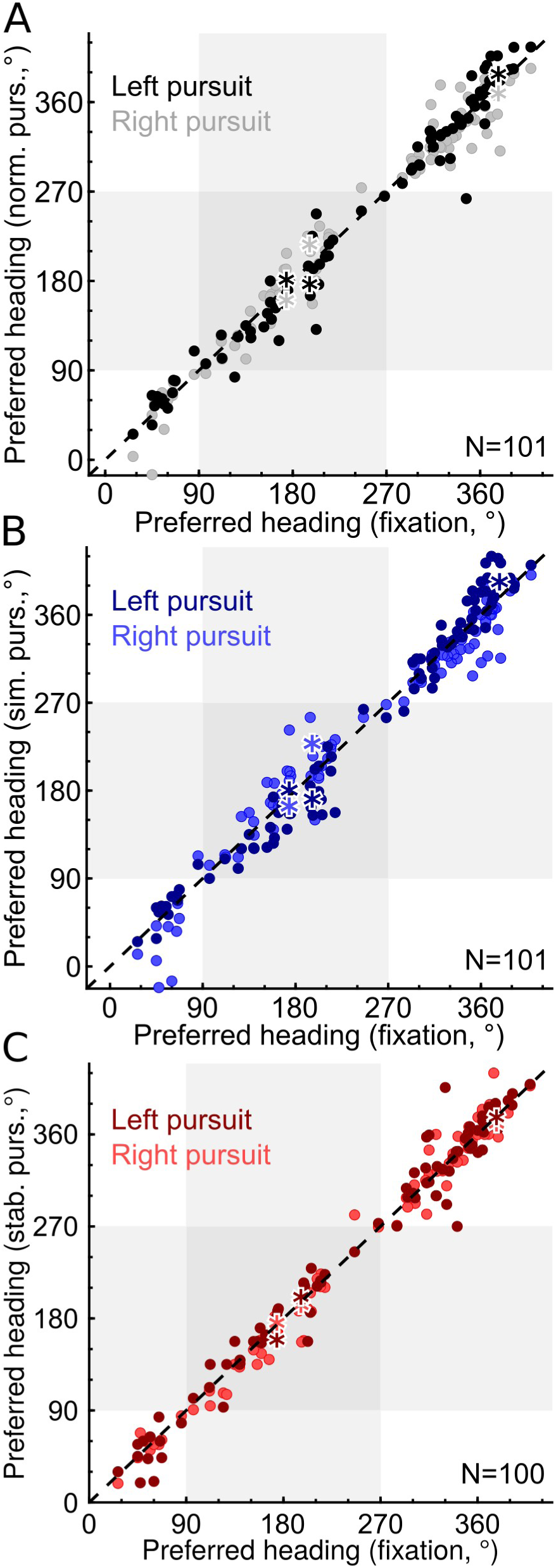
Changes in heading preferences across the cell sample under different pursuit manipulation conditions. Shaded regions denote headings in the backward-going half of heading space. Asterisks mark example cells shown in Figure 4. **A**, Scatter plot relating best fit heading direction preferences under the fixation and normal pursuit conditions. **B**, As in **A**, comparing fixation and simulated pursuit. **C**, As in A and B, but comparing fixation and stabilized pursuit.

In simulated pursuit (Figure 6B), a similar pattern is seen, but with greater departures from the unity line. This result is consistent with the example cells in Figure 4B&D, showing that retinal mechanisms alone are insufficient to maintain stable responses during smooth pursuit. In stabilized pursuit (Figure 6C), the effects are smaller (more closely clustered around the unity line), but opposite in sign from the results of simulated pursuit (Figure 6B). This result is consistent with the sign of the prediction from the efference-copy hypothesis, but suggests that these signals by themselves have only very subtle effects on the tuning of MSTd neurons.

To examine the magnitudes of these changes more closely, we represent the same data as sample histograms in Figure 7. We find significant shifts in heading tuning for the normal pursuit versus fixation comparison (median = −5.29, p=1.2×10^−5^ Wilcoxon signed-rank test) and for the simulated pursuit versus fixation comparison (median = −8.64, p=9.02×10^−15^ Wilcoxon signed-rank test). The stabilized pursuit versus fixation comparison shows a small shift in the direction opposite of the normal pursuit comparison, but this is not significant (median = 0.751, p = 0.0596 Wilcoxon signed-rank test). Overall, we find that the shift in heading preference is only a modestly larger during simulated pursuit than during normal pursuit, and no significant tuning shifts occur during stabilized pursuit. There were no significant differences in results among the three animals (ANOVA; Table 3). Taken together, our results are inconsistent with the hypothesis that efference copy alone can substantially alter heading tuning in MSTd during smooth pursuit.

**Table 3.** Results from 3-way ANOVA for inter-animal differences in tuning shifts.

|  | Normal vs. Fixation | Simulated vs. Fixation | Stabilized vs. Fixation |
| --- | --- | --- | --- |
| Monkey | F = 0.156, p = 0.855,<br>df = 2 | F = 0.553, p = 0.576,<br>df = 2 | F = 2.03, p = 0.134,<br>df = 2 |
| Pursuit Direction | F = 2.15, p = 0.144,<br>df = 1 | F = 0.411, p = 0.522,<br>df = 1 | F = 1.03, p = 0.312,<br>df = 1 |
| Heading Direction | F = 0.997, p = 0.319,<br>df = 1 | F = 1.98, p = 0.161,<br>df = 1 | F = 0.570, p = 0.451,<br>df = 1 |
| Monkey*Pursuit Dir. | F = 1.01, p = 0.368,<br>df = 2 | F = 1.34, p = 0.264,<br>df = 2 | F = 0.054, p = 0.947,<br>df = 2 |
| Monkey*Heading Dir. | F = 1.12, p = 0.328,<br>df = 2 | F = 1.74, p = 0.178,<br>df = 2 | F = 0.178, p = 0.837,<br>df = 2 |
| Pursuit*Heading Dir. | F = 0.529, p = 0.468,<br>df = 1 | F = 1.62, p = 0.205,<br>df = 1 | F = 0.163, p = 0.687,<br>df = 1 |

A 3-way ANOVA was run individually for each of the three different pursuit conditions. Monkey ID, Pursuit Direction, and Heading Direction are included as factors, with the interaction between below these effects.

**Figure 7.**
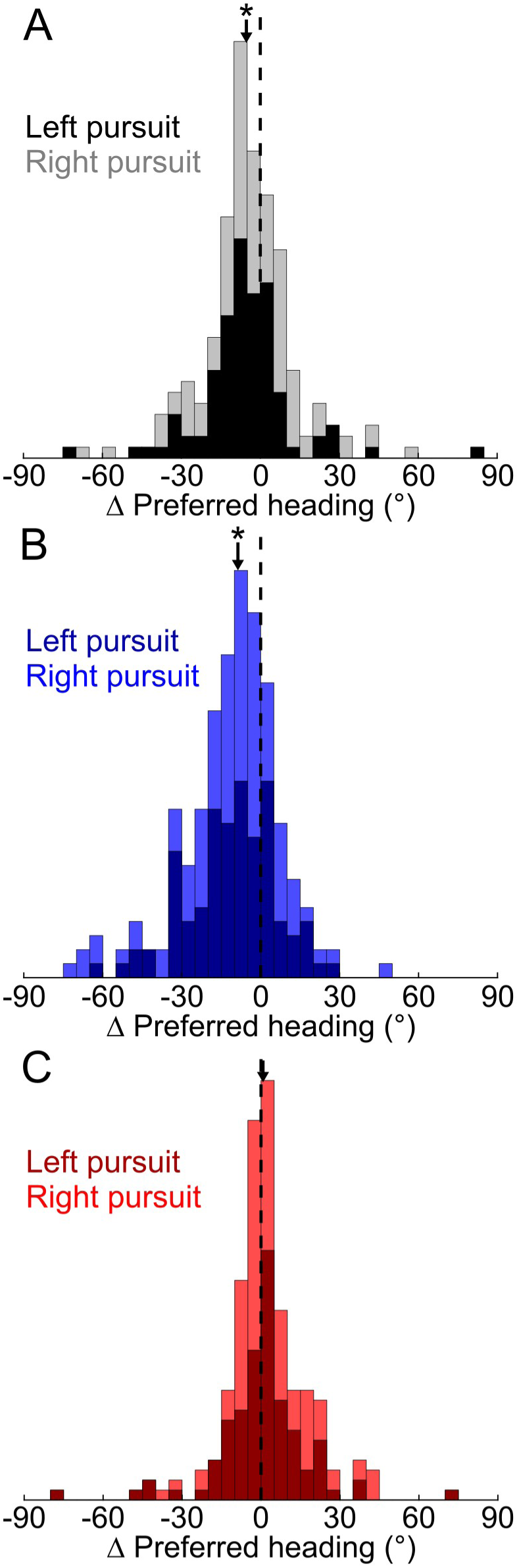
Changes in heading preferences across entire cell sample under different pursuit manipulation conditions. To produce a consistent expected direction of tuning shift (i.e. forward heading and rightward pursuit), the sign of the change was reversed for cells tested with backward headings and leftward pursuit (see Methods). **A**, Normal pursuit. **B**, Simulated pursuit. **C**, Stabilized pursuit. Significance for shift in preferred heading direction between fixation and each pursuit condition was determined with a Wilcoxon Signed-Rank test and threshold of p<0.05. Median paired differences are marked with black arrow and significant differences with asterisks.

### Extraretinal signals have a relatively small influence on heading tuning in MSTd neurons compared to retinally based compensatory signals

Because individual cells in the sample differed in their shift magnitudes, we were interested in whether these differences were systematic or random. If MSTd neurons achieve pursuit stability solely through an efference-copy mechanism, then the shift in heading preference between the normal-simulated and stabilized pursuit-fixation pairs should be of equal magnitude and sign (Figure 3B). Thus for forward heading and rightward pursuit, efference copy would shift cell heading preferences rightward compared to conditions without efference copy. We reasoned that the cells that showed larger shifts during stabilized pursuit should also show larger shift differences between normal and simulated pursuit, as they rely more on efference-copy mechanisms and less on retinal mechanisms. To test this, we first directly compared the shifts between normal and simulated pursuit on a cell-by-cell basis (Figure 8A), using the same folding transform as in Figure 7. We find that heading preferences shifted significantly more during simulated pursuit than during normal pursuit (median = 3.14, p = 3.02×10^−5^, Wilcoxon signed-rank test). When we compare these differences with the shifts during stabilized pursuit (Figure 8B), we find that there is a significant positive correlation between the two (r = 0.387, p = 1.15×10^−7^). A model II regression found that the paired magnitudes of the two shifts are nearly equal (slope = 0.885). Using a permutation test (see Methods), we identified significant shift differences between normal and simulated pursuit (N=13), stabilized pursuit and fixation (N=11), or both (N=2). In sum, these results are consistent with a small but significant contribution of efference copy to heading preferences during pursuit for a minority (22 of 101) of cells.

The small magnitude of extraretinal contributions is somewhat surprising given the substantial distortions to optic flow during pursuit and the larger predictions made in the psychophysical literature (Royden et al., 1994; Banks et al., 1996). Therefore, we investigated the magnitude of correction resulting from a retinal mechanism in MSTd neuronal responses. The contribution of these corrections should appear as the difference between the hypothetical tuning shift for a completely uncorrected neuron’s responses (which would be located to the left of the simulated curves in Figure 3B) and the cell’s shift seen in simulated pursuit. To calculate this uncorrected tuning shift, we first needed to estimate how MSTd neurons respond to the apparent path curvature in distorted retinal flow patterns. When the eyes are stationary during forward self-motion, the centers of motion in each depth plane are all at the same retinal location, which indicates the heading direction. During pursuit, however, the centers will shift on the retina in the direction of eye movement (or in the opposite direction for backward self-motion). The magnitude of this shift is proportional to the distance of the plane from the observer (Figure 9A), producing curvature in the flow pattern. Psychophysical evidence finds that observers perceive a self-motion direction in curved flow patterns that is coincident with the closest visible depth planes in the stimulus (Royden, 1994). Physiological evidence also suggests that near depth planes should dominate MSTd responses, as these neurons generally prefer higher speeds, and motion parallax produces higher speeds at close distances (Tanaka and Saito, 1989; Upadhyay et al., 2000; Inaba and Kawano, 2010).

**Figure 9.**
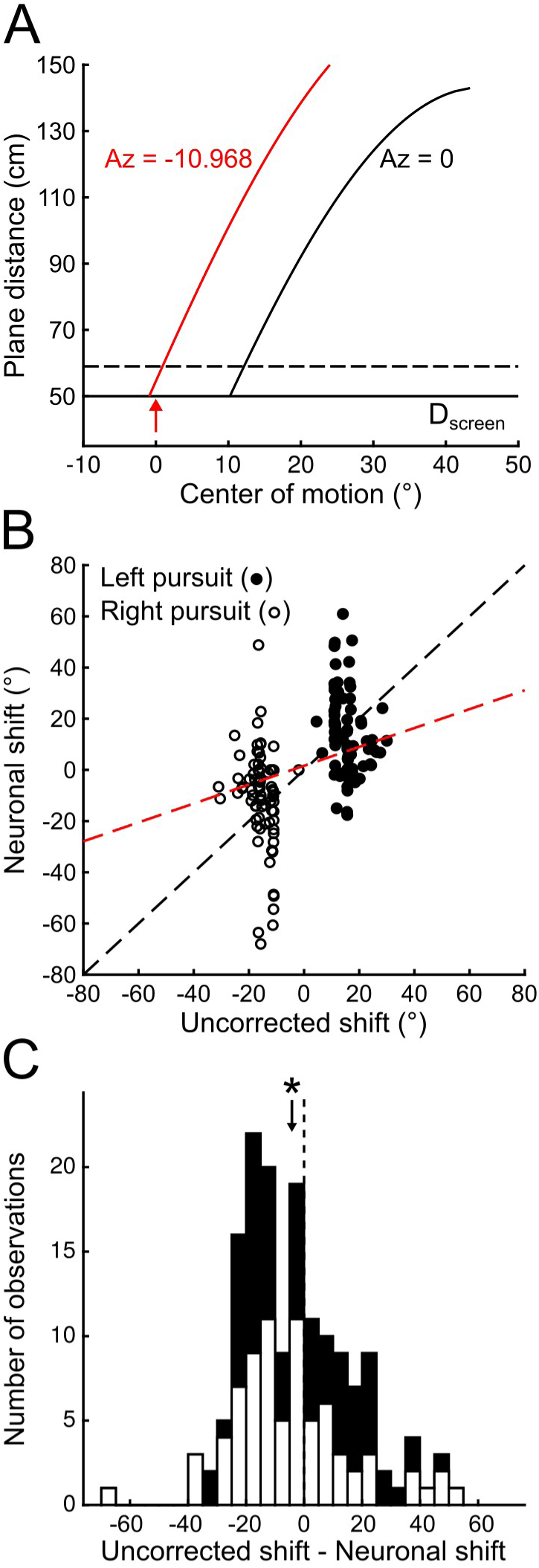
Effects of retinally based correction on heading tuning in our sample. **A**, Method of calculating center of motion shifts on the retina. For each cell’s preferred heading direction (0°Az, here), the retinal position of the center of motion during rightward pursuit at 10°/s was calculated for each depth plane in the stimulus (black curve; see Methods). Under these conditions, the heading direction that produces a mean center of motion at 0° across the nearest 10 cm of the stimulus (red arrow) is about −11°. **B**, Comparison scatter plot of the calculated uncorrected shifts and the neuronal shifts (preferred heading in simulated pursuit - fixation). Line of best fit calculated with linear regression. **C**, Paired comparison of uncorrected and neuronal shifts. The sign shifts for leftward pursuit have again been flipped to be consistent with rightward pursuit. Median paired difference between uncorrected and neuronal shifts is shown with a black arrow, and significance is denoted with an asterisk (Wilcoxon Signed-Rank test).

Therefore, to build a set of uncorrected responses, we assigned each retinal flow pattern presented during simulated pursuit a heading direction based on the mean center of motion in the nearest depth planes of the stimulus. This procedure is outlined in Figure 9A for a neuron that prefers a forward heading direction (0°Az) during fixation. During rightward simulated pursuit, the retinotopic locations of the centers of motion during pursuit are shifted by about 10° to almost 50° of visual angle (curved black line) away from their preferred retinal location. We estimate the heading direction for a MSTd neuron that did not compensate for pursuit in this case by finding the average location of the centers of motion for the nearest 10 cm of the stimulus (i.e. between the full and dashed horizontal black lines). For this neuron tuned to 0°, this is clearly a sub-optimal stimulus.

To find the optimal stimulus, we can instead find a heading direction (see Methods) that produces a mean center of motion during pursuit at the cell’s preferred location (0°). Our expected *uncorrected shift* in heading preference for the example neuron in Figure 9A is obtained by the difference between this heading and the cell’s preferred heading during fixation (∼ −11° in this example). We ran this procedure across all neurons and compared the calculated uncorrected shift and the actual shifts seen during simulated pursuit (Figure 9B). Across our sample and both pursuit directions, a linear regression shows that the two are related with a best fit line of slope = 0.369. The variation in the calculated uncorrected shifts is due to a slightly positive relationship between flow curvature and preferred heading laterality (e.g., Figure 9A). After sign-reversing the data to be consistent with forward self-motion and rightward pursuit, we find that the uncorrected shift is significantly more negative than the measured neuronal shift in simulated pursuit (Figure 9C; median = −5.22°, p = 0.012). These results form an upper bound on the magnitude of retinally based compensation, given that they are based on the depth planes with the smallest magnitude of center of motion shifts on the retina. If instead we take a weighted mean over all planes (either equally weighted or weighted inversely by the distance from the observer), the paired difference is much more substantial (unweighted: median shift = −18.6, p =1.41×10^−12^; weighted: median shift = −13.1, p = 6.22×10^−10^). Taken together, we find a significant contribution of a retinal mechanism to the stability of MSTd neurons during pursuit that is larger than that of the efference-copy contribution.

### Viewing condition and pursuit accuracy do not significantly alter heading preference changes

We wanted to confirm that our inclusion of two different viewing conditions did not significantly influence our results. To do so, we recorded from a subset of cells under both binocular and monocular viewing, when cells could be held for long enough. A paired comparison (Figure 10) failed to show a significant median difference in heading preference shifts between the two viewing conditions (normal vs. fixation: median = 0.219, p = 0.411; simulated vs. fixation: median = 0.662, p = 0.89; stabilized pursuit vs. fixation: median = −1.01, p = 0.42, Wilcoxon signed-rank test).

We also investigated whether differences in eye movements between each of the pursuit conditions could account for the changes in heading preference. During fixation and simulated pursuit, the pursuit gain is fixed at one (i.e. the mean horizontal eye velocity in a given trial is equal to the horizontal velocity of the pursuit target), while in normal and stabilized pursuit it depends on each animal’s performance. This result could inflate the shift difference between normal and simulated pursuit if eye speed is substantially lower than that of the pursuit target. The mean pursuit gain across all trials with active pursuit and across the three monkeys is 0.88 ± 0.069 (standard deviation), which is similar to previous reports for tracking eye movements in the presence of a textured background (Takeichi et al., 2003).

While this value is below unity, gains for each condition were quite variable. We took advantage of this variability to see if pursuit gain predicted the magnitude of shifts in heading preference. We found no significant correlation between average pursuit gain and heading preference shift on a cell-by-cell basis (normal pursuit, left: r = 0.0735, p = 0.50, right: r = 0.0022, p = 0.984; stabilized pursuit, left: r = −0.0013, p = 0.991, right: r = 0.114, p = 0.310). In a three-way ANOVA across all animals and trials with pursuit condition (normal or stabilized pursuit), pursuit direction, and heading laterality (within or outside 45° forward or backward headings) as factors, we found a significant main effect for pursuit direction (F = 77.09, p = 7.30×10^−17^, df = 1). The other factors did not significantly account for gain variance (pursuit condition: F = 2.82, p = 0.094, df = 1; heading laterality: F = 2.9, p = 0.089, df = 1). Right pursuit had higher mean gain than left in both normal and stabilized pursuit (mean difference of 0.056 and 0.06 respectively, or 6.4% and 6.8% of the overall mean gain). This result was likely due to a slight bias in our sample for neurons preferring leftward headings, which caused us to present more leftward headings in the main experiment. Lateral heading directions produce flow patterns that drive optokinetic nystagmus (Miles, 1995); a bias for one direction of flow over the other would produce a selective reduction in pursuit gain in one direction (and an increased gain in the other direction) via the interaction between the slow phase of optokinetic nystagmus and smooth pursuit. In the end, however, this pursuit gain effect was small and does not appear to have a significant effect on tuning shifts given the results of the pursuit direction factor in Table 3.

We also tested for significant differences in the number of saccades between each of the pursuit conditions. Though we controlled for initial catch-up saccades as much as possible with a step-ramp pursuit stimulus (see Methods), full-field visual motion may produce optokinetic nystagmus that has a saccade-like fast phase. The mean number of saccades across all pursuit conditions was 1.84 ± 0.616 (standard deviation). A three-way ANOVA with the same three factors (including fixation and simulated pursuit as components of the pursuit condition factor) reveals significant effects of each on the mean number of saccades (pursuit condition: F = 106.1, p = 5.58×10^−41^, df = 2; pursuit direction: F = 11.61, p = 6.95x10^−4^, df = 1; heading laterality: F = 5.52, p = 0.0191, df = 1). The largest differences were between conditions in which the eyes were fixated (fixation and simulated pursuit) versus active-pursuit conditions (normal and stabilized pursuit). This finding was likely due to an interaction between smooth pursuit and the fast phase of optokinetic nystagmus, as pursuit eye movements tend to suppress optokinetic nystagmus (Post et al., 1984). In absolute numbers, however, these inter-condition differences were small (maximum absolute mean paired difference = 0.704 saccades/trial).

### Pursuit manipulations do not substantially affect other aspects of neuronal response patterns

Our analysis thus far has focused on the stability of heading tuning preferences during pursuit eye movements, but theoretical work has proposed that MSTd can encode veridical heading direction at the population level with gain fields or by modulating response offset. Gain fields for eye velocity may reflect the first stage of the process used to produce shifts in heading preference, as found for *positional* coordinate transforms in parietal areas (Andersen et al., 1985; Beintema and van den Berg, 1998). Response offsets (i.e., the response of the cell to the antipreferred heading) of the cell population could be additively modulated with an efference-copy signal to perform a vector operation that would subtract off the pursuit-related components from retinal flow (Perrone and Krauzlis, 2008).

To test these hypotheses, we investigated whether other parameters of the cell-response fits were modulated between fixation and the different pursuit conditions. These parameters (Figure 4A, bottom) were extracted at the same time as the estimation of each cell’s preferred direction using a modified von Mises function in which all parameters are independent (see Methods). Overall, the cells showed a decrease in their firing range across all three pursuit conditions, as seen in the response-amplitude parameter (Figure 11; normal vs. fixation: median = −3.29, p = 2.91×10^−5^; simulated vs. fixation: median = −1.75, p = 1.87×10^−4^; stabilized pursuit vs. fixation: median = −2.75, p = 6.16×10^−6^, Wilcoxon Signed-Rank). Neither the tuning bandwidth (normal vs. fixation: median = −1.17, p =0.593; simulated vs. fixation: median = −4.53, p = 0.179; stabilized pursuit vs. fixation: median = −2.08, p = 0.0823, Wilcoxon Signed-Rank), nor the offset (normal vs. fixation: median = −2.42×10^−7^, p = 0.693; simulated vs. fixation: median = −1.15×10^−6^, p = 0.404; stabilized pursuit vs. fixation: median = −2.37×10^−7^, p = 0.669, see Methods for permutation test) parameters showed any significant difference between fixation and pursuit across the cell sample. Further analysis revealed no significant differences in parameter changes between left and right pursuit (Table 4).

**Table 4.** Differences in tuning curve parameters between left and right pursuit.

|  | Normal Pursuit (L vs. R) | Simulated Pursuit (L vs. R) | Stabilized Pursuit (L vs. R) |
| --- | --- | --- | --- |
| Response Amplitude | Median diff. = 0.945sp/s,<br>$p = 0.179$ | Median diff. = 1.62sp/s,<br>$p = 0.221$ | Median diff. = 0.393sp/s,<br>$p = 0.504$ |
| Response Offset | Median diff. = -1.03sp/s,<br>$p = 0.726$ | Median diff. = -3.48sp/s,<br>$p = 0.617$ | Median diff. = -1.45sp/s,<br>$p = 0.543$ |
| Tuning Bandwidth | Median diff. = $-6.22^\circ$,<br>$p = 0.304$ | Median diff. = $4.75^\circ$,<br>$p = 0.208$ | Median diff. = $-1.39^\circ$,<br>$p = 0.969$ |

Paired comparisons between tuning parameter fits for left and right pursuit. Wilcoxon signed-rank test.

**Figure 11.**
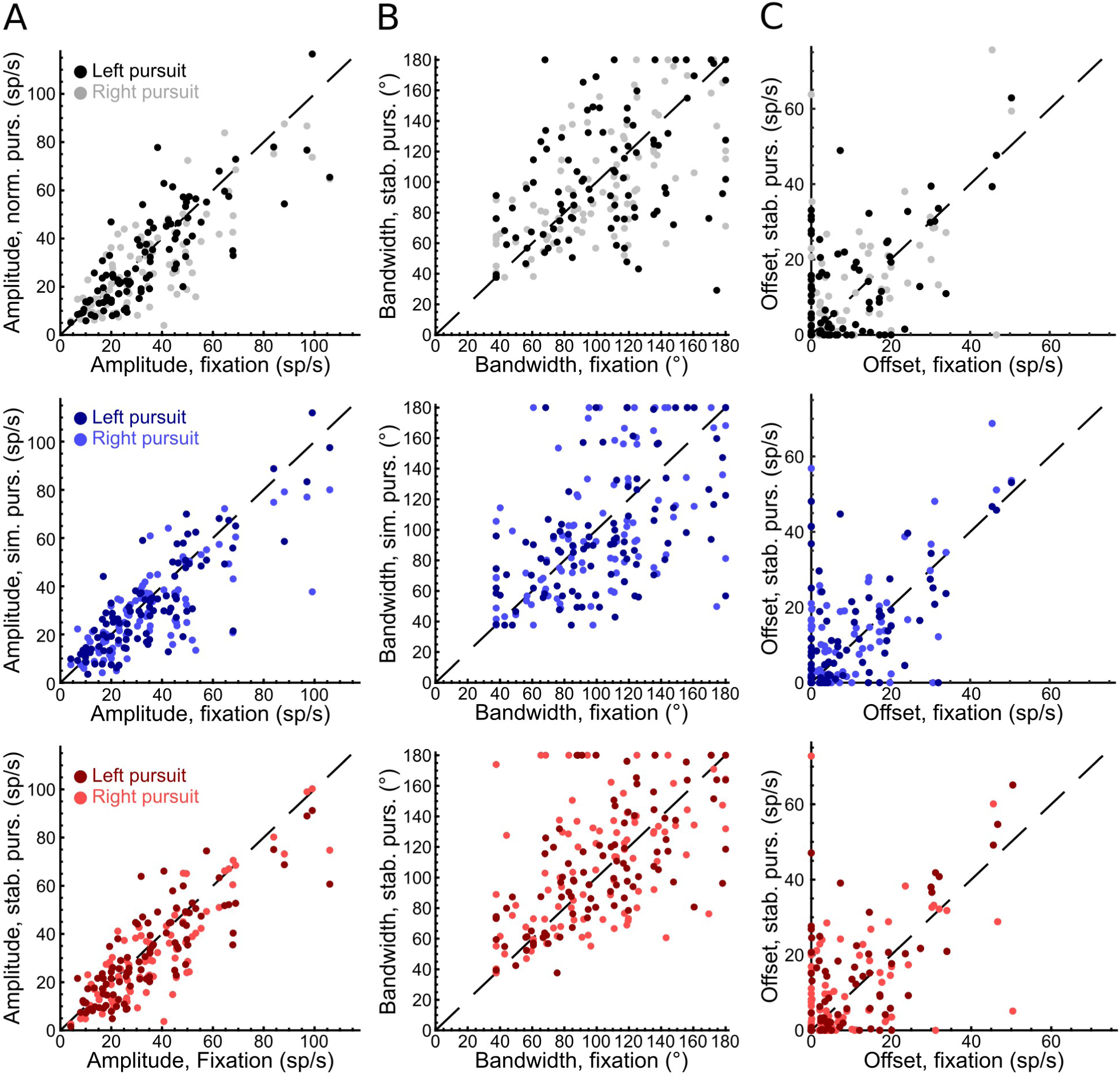
Effects of pursuit manipulations on other tuning curve parameters. Each plot is a cell-by-cell paired comparison of best fit parameters for fixation and pursuit. **A**, Amplitude parameter. **B**, Tuning curve bandwidth or FWHM, as derived from sigma parameter. **C**, Response offset parameter (sometimes referred to as baseline).

Offset parameters were most often best fit with a value of zero, which taken together with the median change across the sample is at odds with theoretical mechanisms that depend on this response property (Perrone and Krauzlis, 2008). Although we were not able to replicate previous findings of gain fields for eye velocity (Squatrito and Maioli, 1997) with the amplitude parameter, our sampling of pursuit space was limited to two different directions that were often misaligned with the cells’ null-preferred pursuit direction axes. In sum, we find that our pursuit manipulations most directly affected neurons’ preferred heading directions rather than other aspects of cell tuning.

## Discussion

We investigated how animals compensate for the sensory consequences of their own behavior, using monkey visual-motion perception as a model. Our experiments measured the relative contributions of extraretinal and retinal mechanisms to the stability of heading responses in MSTd during smooth pursuit. To this end, we developed a novel manipulation of an optic flow stimulus that actively stabilizes the flow pattern on the retina during smooth pursuit to eliminate the effects of retinal mechanisms on the heading preferences of MSTd neurons. Using this manipulation alongside a simulated pursuit paradigm revealed that the contributions of extraretinal mechanisms to the stability of heading tuning during pursuit are small compared to the contributions of retinal mechanisms. These results show that high-level visual cortex discounts incoming sensory reafference mainly through retinal mechanisms when depth cues are available.

### Relationship with previous physiological work

Our work differs in some important ways from previous experiments that identified a larger role for efference copy in MSTd. The Andersen lab (Bradley et al., 1996; Shenoy et al., 2002) reported large changes in preferred headings between normal and simulated pursuit, which could be due to a variety of factors. Their flow stimulus included only a single depth plane, was limited to a 50° by 50° field of view, and had a higher ratio of pursuit speed to self-motion speed (which produces a larger set of center of motion shifts in the pattern).

Most proposed retinal mechanisms of pursuit compensation depend on the depth structure of the scene. Because the components of retinal flow due to retinal slip affect all depth planes equally, while the self-motion components are subject to motion parallax, the brain could theoretically subtract the full-field slip to retrieve the undistorted pattern of motion (Longuet-Higgins and Prazdny, 1980; Heeger and Jepson, 1992; Royden, 1997(Longuet-Higgins and Prazdny, 1980; Heeger and Jepson, 1992; Royden, 1997)). With only a single depth plane available, MSTd neurons would depend entirely on efference copy to solve the rotation problem, increasing the difference in heading preference shifts between normal and simulated pursuit due to increased shifts during simulated pursuit (as discussed above; see Figure 9). Thus the single-plane stimulus enables an accurate and assumption-free estimate of the retinal shift, but at the cost of removing the most profound cue that enables a retinally based solution.

Conclusions similar to ours have been drawn about the dominance of retinal stability mechanisms in ventral intraparietal sulcus (VIP) (Sunkara et al., 2015). VIP represents heading direction at a nearly identical level of precision to MSTd and is similarly tolerant to pursuit distortions (Maciokas and Britten, 2010). It also appears to code heading direction gleaned from optic flow in eye-centered coordinates like MSTd (Chen et al., 2013), though some neurons’ receptive fields are more consistent with a head-centered coordinate system (Duhamel et al., 1997 (Duhamel et al., 1997)). Sunkara and colleagues also find only small differences in preferred heading shifts between normal and simulated pursuit, and conclude retinal factors contribute more to pursuit tolerance in neurons than does efference copy. Their study differs from ours and those previously mentioned in their inclusion of binocular depth cues, which provide additional information for a retinally based stability mechanism. Changes in the disparity structure in the scene can have profound psychophysical effects on apparent shifts in heading direction during simulated pursuit (van den Berg and Brenner, 1994; Grigo and Lappe, 1998). Both MT and MSTd neurons are sensitive to disparity and may use this cue instead of efference copy to compute true heading direction during pursuit (Kim et al., 2015). As in our current study, however, Sunkara and colleagues do not find substantial differences in tuning shifts between monocular and binocular viewing conditions during simulated and normal pursuit. They propose that motion parallax and dynamic perspective (i.e., the changing angle between the viewing direction and the visual scene during pursuit) cues alone can support retinally based mechanisms. Overall, the richness of these retinal cues helps explain why our results show that retinal mechanisms dominate pursuit compensation in higher-level visual cortex.

### Theoretical retinal mechanisms

How neurons use retinal cues to extract the true heading from the distorted flow patterns is debated. One group proposed that MSTd neurons directly encode heading direction with a dense array of templates that combine varying amounts of retinal slip with optic flow fields resulting from self-motion (Perrone and Stone, 1994, 1998). A downstream decoder would then choose the preference of most active neuron as the current heading direction. Others suggested that the true heading is instead extracted at the population level via sparse decomposition of the flow field (Beyeler et al., 2016). The resulting units were MSTd-like with preferences for radial, rotating, and laminar motion, but with patchier spatial extents of their receptive fields. Subsequent decoding of the population with a linear combination of units could recapitulate some (but not all) of the psychophysical results comparing heading biases between simulated and normal pursuit.

Alternatively, other groups proposed retinal slip is separated from flow due to self-motion by combining tuning shifts with gain modulation of neuronal responses (Beintema and van den Berg, 1998). Analysis of single-unit responses in VIP to flow patterns with different combinations of translational and rotational components revealed that some cells showed joint, inseparable tuning for specific combinations (Sunkara et al., 2016). These responses were similar to those seen in real pursuit, suggesting that VIP simultaneously represents both the distorted retinal flow pattern and the veridical heading, which could be extracted by a downstream area. Similar joint tuning may be present in MSTd. Along this line, another group also separately extracted pursuit direction and the center of motion in head-centered coordinate using an optimal linear estimator on MSTd responses to optic flow from a single depth plane, concluding that MSTd encodes eye and self-motion with a set of basis functions (Ben Hamed et al., 2003).

These different model formulations make testable predictions for future experiments. The spatial extent of MSTd receptive fields can be revealed by further investigation into responses to local motion patches and interactions between patches (Heuer and Britten, 2007; Mineault et al., 2012). Although our stimulus paradigm was not designed to test these models directly, one could identify joint tuning curves for pursuit and heading with denser sampling of pursuit space within the simulated pursuit paradigm.

### Relationship with human psychophysical literature

Despite our physiological findings in MSTd, many psychophysical results support the hypothesis that pursuit compensation in heading perception depends on efference-copy signals (Royden et al., 1994; Banks et al., 1996; Haarmeier et al., 1997). These results may be sensitive to the exact parameters of the experiment, including pursuit speed (Warren and Hannon, 1988), task instructions (Li et al., 2006), and the presence of reference objects in the visual scene (Li and Warren, 2000). Our animals were not trained to make heading judgments, which may have affected top-down influences on sensory cortex. The pursuit and self-motion speeds used in our experiments were similar to those used in Royden et al. 1994. Our stimulus also covered a larger portion of the visual field and contained more dots than theirs, both of which provided richer cues for a retinal mechanism. The size of the field of view especially has been linked to the precision with which observers can identify the axis of eye rotation and the center of motion in retinal flow (Koenderink and van Doorn, 1987). Going forward, it will be critical to identify how retinal and extraretinal mechanisms might be weighted differently at the neuronal level based on these parameters. Some conditions (e.g., scenes with limited depth cues) may force heading-processing areas to rely more on efference copy, while the rich retinal inputs may allow retinal mechanisms to dominate when they are available (Crowell and Andersen, 2001; Wilkie and Wann, 2002).

To resolve these differences between our results and those from the human literature, the stabilized pursuit manipulation needs to be used in psychophysical experiments. We predict that biases in this manipulation would be small, as we see in the physiology. If, however, biases remain as large as they are under simulated pursuit, that finding would suggest that downstream areas integrate efference-copy cues. Relatively few physiological studies have investigated pursuit compensation in frontal (Yang and Gu, 2017), premotor (Cottereau et al., 2017), or high-level parietal (Siegel and Read, 1997) cortical areas that receive projections from high-level visual-motion cortex. Additional investigations into the roles of efference copy in these areas would therefore benefit the field.

## Acknowledgments

National Eye Institute Grant R01 EY022087 (KHB)

National Eye Institute Training Grant T32 EY015387

National Eye Institute Core Facilities Grant P30 EY012576

New Zealand Marsden Fund Grant (JP)

We thank Daniel Sperka for design of code for stimulus display, data recording, and image stabilization. We thank Martin Banks, John Perrone, and Richard Krauzlis for helpful discussion and Conor Weatherford for technical support and animal care.

